# HAPPILEE: The Harvard Automated Processing Pipeline In Low Electrode Electroencephalography, a standardized software for low density EEG and ERP data

**DOI:** 10.1101/2021.07.02.450940

**Authors:** K.L. Lopez, A.D. Monachino, S. Morales, S.C. Leach, M.E. Bowers, L.J. Gabard-Durnam

## Abstract

Low-density Electroencephalography (EEG) recordings (e.g. fewer than 32 electrodes) are widely-used in research and clinical practice and enable scalable brain function measurement across a variety of settings and populations. Though a number of automated pipelines have recently been proposed to standardize and optimize EEG preprocessing for high-density systems with state-of-the-art methods, few solutions have emerged that are compatible with low-density systems. However, low-density data often include long recording times and/or large sample sizes that would benefit from similar standardization and automation with contemporary methods. To address this need, we propose the HAPPE In Low Electrode Electroencephalography (HAPPILEE) pipeline as a standardized, automated pipeline optimized for EEG recordings with low density channel layouts of any size. HAPPILEE processes task-free (e.g. resting-state) and task-related EEG, and event-related potential (ERP) data, from raw files through a series of processing steps including filtering, line noise reduction, bad channel detection, artifact rejection from continuous data, segmentation, and bad segment rejection that have all been optimized for low density data. HAPPILEE also includes post-processing reports of data and pipeline quality metrics to facilitate the evaluation and reporting of data quality and processing-related changes to the data in a standardized manner. We describe multiple approaches with both recorded and simulated EEG data to optimize and validate pipeline performance. The HAPPILEE pipeline is freely available as part of HAPPE 2.0 software under the terms of the GNU General Public License at: https://github.com/PINE-Lab/HAPPE.

## Introduction

Electroencephalography (EEG) recordings are a useful and noninvasive tool for interrogating human brain function across the lifespan. Advancements in the field have allowed for rich data collection in laboratories through the use of high-density channel layouts, but it is not always feasible or optimal to rely on these dense layouts across settings. Low-density channel layouts (fewer than 32 channels) continue to be heavily used, particularly with clinical populations, both in clinical research (Brito et al., 2019, 2016; Gu et al., 2018; Pellinen et al., 2020; van den Munckhof et al., 2018) and diagnostic testing (Aeby et al., 2021; Cassani et al., 2017; Paul et al., 2019; Tiwari et al., 2017), as well as in low-resource areas (Kariuki et al., 2016; Siddiqi et al., 2015; Sokolov et al., 2020; Williams et al., 2019). A low-density EEG approach also provides the flexibility for researchers to travel to participants for testing in natural contexts (e.g. school-based or home-based studies, Troller-Renfree et al., 2021) or in the event that participants cannot come to the lab. Low-density EEG will also be instrumental in future research, given the current momentum towards large-scale neuroscience studies that achieve community implementation and the focus on precision medicine through brain-based biomarkers (e.g. potential EEG-based screening for Autism Spectrum Disorder at well-child doctor’s visits), where high-density recordings may be neither practical nor necessary. Indeed, a number of wearable, ultra-low-cost, low-density EEG hardware solutions are emerging in industry to facilitate such measurement. A key impediment to the use of low-density EEG in these contexts is the fact that the raw EEG signal is contaminated by both environmental and physiological artifacts. Up to this point, researcher selection of uncontaminated EEG data has been the field’s standard, but even with low-density data, this method is time-consuming, subjective, and does not allow for the efficient processing of a large number of data sets. Low-density EEG collected in clinical contexts that can span hours may also preclude manual inspection due to recording length. As a result, there remains a current and growing need for software that standardizes and automates the processing and removal of artifacts in low-density EEG data.

There is now an extensive collection of automated EEG processing pipelines (e.g., Andersen, 2018; APP, da Cruz et al., 2018; MADE, Debnath et al., 2020; EEG-IP-L, Desjardins et al., 2021; HAPPE, Gabard-Durnam et al., 2018; Hatz et al., 2015; FASTER, Nolan et al., 2010; Automagic, Pedroni et al., 2019; EPOS, Rodrigues et al., 2020). However, their reliance on independent component analysis (ICA) to segregate and reject artifacts makes them unsustainable for low-density data, as the limited number of channels provides insufficient independent components for robust artifact isolation. Many of these pipelines also use standard deviation-dependent approaches to identify and reject outlier data or channels as artifact-contaminated. These approaches may require modification to scale down to low-density setups with few channels. Other software tools built into these fully-automated pipelines to aid in different stages of artifact detection and rejection are most effective when used with high-density data or have not been validated in low-density data (PREP, Bigdely-Shamlo et al., 2015; SASICA, Chaumon et al., 2015; Adjusted-ADJUST, Leach et al., 2020; ADJUST, Mognon et al., 2011; ASR, Mullen et al., 2013; MARA, Winkler et al., 2014). A recent automated mega-analysis by Bigdely-Shamlo et al., 2020 introduced a pipeline that supports both high-density and low-density data, but only at the upper bound of low-density channel layouts. Specifically, they tested their pipeline using data sets ranging from a Neuroscan 30-channel headset to a Biosemi 256-channel headset, but found that the density of the headset accounted for variability in the channel amplitudes across datasets after processing. Several pipelines automate the processing of strictly low-density data. Of these options, some are made specifically for a particular population (Cassani et al., 2017) or acquisition systems (e.g. James Long EEG Analysis System software, Whedon et al., 2020). Others use independent component analysis to reject artifacts (Hajra et al., 2020), which cannot support many low-density setups, or offer only simple artifact-rejection approaches like segment rejection that can cause significant data loss without artifact-rejection in continuous data first (e.g. EEG Analysis System software). Thus, there remains a need for standardized processing solutions to serve the range of low-density EEG configurations in use.

To address this need, we propose a novel pipeline for low-density EEG data called HAPPILEE (Harvard Automated Preprocessing Pipeline Including Low-Electrode Encephalography). We apply contemporary approaches to optimize line noise reduction, bad channel detection, artifact rejection from continuous data, segmentation, and bad segment rejection methods to suit low density datasets. The following sections of this manuscript describe HAPPILEE’s processing steps and outputs, assess optimization of these steps for low-density EEG/ERP data, and demonstrate HAPPILEE’s effectiveness with a low-density developmental EEG dataset.

### Optimization Dataset

The various steps of the HAPPILEE automated pipeline were optimized using a subset of developmental EEG files from the Bucharest Early Intervention Project (BEIP) (for full study design, see Zeanah et al., 2003). The EEG files contributing to this example dataset may be freely assessed at Lopez et al. on Zenodo. We selected the BEIP dataset as its study design facilitated testing HAPPILEE on EEG data from children across a range of caregiving conditions, behavioral/clinical phenotypes, and ages (Zeanah et al., 2009). The optimization dataset includes resting-state EEG from three groups of children living in Romania starting in 2001. The first group, referred to as the Care as Usual Group (CAUG), is composed of children living across six institutionalized care facilities throughout Bucharest, Romania. The second group is the Foster Care Group (FCG), which is composed of children who were removed from these institutions through random assignment and placed in a foster care intervention. The final group is the Never Institutionalized Group (NIG), made up of a community sample of children living with their biological families who have never been placed in institutionalized care or foster care. We selected a subset of thirty EEG files across the three groups from the greater dataset (CAUG, *n*=8; FCG, *n*=8, NIG, *n*=14). The average age at the start of the baseline assessment across the three groups was 17.40 months, with a range of 6.28 - 29.98 months (averages per group: CAUG=18.30; FCG=16.21; NIG=17.58). The resting-state EEG for all children was recorded with the James Long system from twelve scalp sites (F3, F4, Fz, C3, C4, P3, P4, Pz, O1, O2, T7 and T8) using a lycra Electro-Cap (Electro-Cap International Inc., Eaton, OH) with sewn-in tin electrodes.

### HAPPILEE Data Inputs

HAPPILEE accommodates multiple types of EEG files with different acquisition layouts as inputs. A single run will support only a single file type across files, specified by the user. For .set formatted files, the correct channel locations should be pre-set and embedded in the file (e.g. by loading it into EEGLAB and confirming the correct locations) prior to running through HAPPILEE. When running .mat formatted files, you must have a file with channel locations specified in your folder in order to run all steps in the HAPPILEE pipeline. If channel locations are not provided, you will not be able to do the following: filter to channels of interest, interpolate bad channels, and re-reference your data. Additionally, the data QC output table will not list which bad channels were rejected during the optional bad channel detection step if you do not provide channel locations. Each batch run of HAPPILEE must include files collected with the same channel layout (company, net type, and electrode number) and paradigm (resting-state or event-related), each of which users must specify for a given run. HAPPILEE processes data collected with any sampling rate, and files within a single run may differ in their individual sampling rates (if this is the case, we strongly recommend selecting the option to resample data to the same frequency to ensure subsequent steps perform comparably across files regardless of original sampling rate).

### Line Noise Removal

HAPPILEE removes electrical noise (e.g., 60 or 50 Hz artifact signal) from EEG through the multi-taper regression approach implemented by the CleanLineNoise program (Mullen, 2012). Multi-taper regression can remove electrical noise without sacrificing or distorting the underlying EEG signal in the nearby frequencies, drawbacks of the notch-filtering approach to line-noise removal (Mitra and Pesaran, 1999). Specifically, HAPPILEE applies the updated version of CleanLine’s multi-taper regression (called CleanLineNoise, implemented in the PREP pipeline) which is more effective at removing line noise than the original CleanLine version present in HAPPE 1.0 software (purportedly a bug fix in the CleanLine code, see Makoto’s pipeline page for unpublished evidence). The legacy CleanLine version from HAPPE 1.0 is available as an option to the user, however the updated version is registered as the default. CleanLineNoise’s multi-taper regression scans for line-noise signal near the user-specified frequency ± 2 Hz, a 4-s window with a 1-s step size and a smoothing tau of 100 during the fast Fourier transform, and a significance threshold of p = 0.01 for sinusoid regression coefficients during electrical noise removal. This process is highly specific to the frequency of electrical noise, which the user can specify to be 60 Hz or 50 Hz. Quality control metrics for the degree and effectiveness of line noise removal are automatically generated in HAPPILEE and discussed in detail as part of the subsequent “Quality Control Metrics” section of this manuscript.

### Filtering

HAPPILEE applies an automatic low-pass filter at 100 Hz prior to artifact rejection for all files. If processing data for ERP analyses, additional filtering as determined by the user occurs after artifact rejection. If processing resting-state EEG or task-based data for time-frequency analyses, the filtering at this stage is a band-pass filter from 1-100 Hz.

### Bad Channel Detection (Optional)

HAPPILEE includes an option to detect channels that do not contribute usable brain data due to high impedances, damage to the electrodes, insufficient scalp contact, and excessive movement or electromyographic (EMG) artifact throughout the recording. Users have the option to run bad channel detection or not (in which case all channels are subjected to subsequent processing steps). Various methods are currently used to detect and remove bad channels across automated pipelines. However, some common automated detection methods used for high-density EEG may not be optimal for low-density EEG without modification, especially those relying heavily on standard deviation-related metrics of activity to detect outlier channels. For example, in HAPPE 1.0 (Gabard-Durnam et al., 2018), bad channel detection is achieved by evaluating the normed joint probability of the average log power from 1 to 125 Hz across the user-specified subset of included channels. Channels whose probability falls more than 3 standard deviations from the mean are removed as bad channels in two iterations of this bad channel detection step. However, removing channels that are three or more standard deviations from the mean activity assumes a normal distribution of channel activities (via the Central Limit Theorem) that we cannot assume with low-density channel numbers (Altman and Bland, 1995). Similarly, the FASTER algorithm (Nolan et al., 2010) used in the MADE pipeline flags channels by measuring each channel’s Hurst exponent, correlation with other channels, and channel variance and standardizing the three values with an absolute Z-score (subject to the same constraints as standard deviations with very small samples). Thus, these algorithms require validation before implementation in low-density EEG data.

Other methods like EEGLab’s Clean Rawdata algorithm may more readily translate to low-density EEG data. Specifically, Clean Rawdata’s ‘Flatline Criterion,’ can detect channels with flat recording lengths longer than a user-specified threshold of seconds (indicating no data collected at that location). If the channel contains a flatline that lasts longer than the threshold, the channel is marked bad. Similarly, ‘Channel Correlation Criterion’ sets the minimally acceptable correlation value between the channel in question and all other channels. If a channel is correlated at less than the preset value to an estimate based on other channels, it is considered abnormal and marked bad. But features like the Line Noise Ratio Criterion, which identifies whether a channel has more line noise relative to neural signal than a predetermined value, in standard deviations based on the total channel population, should be assessed in the low-density EEG context.

To test the efficacy of FASTER, HAPPE 1.0, and Clean Raw data functions and determine the optimal criterion values for the detection of bad channels in low density data, we compared a series of thirty-three automated options to a set of manually identified bad channels for nineteen files in the BEIP dataset (this subset of files had strong agreement about good/bad channel labels by three field experts each with over a decade of EEG experience). Specifically, the files were run through the HAPPE 1.0 legacy detection method for bad channels and the FASTER detection method used in the MADE pipeline, as well as a number of iterations of the Clean Rawdata function and combinations of Clean Rawdata with spectrum evaluation to optimize channel classification (shown in Table 1). Note that for iterations of Clean Rawdata with Flatline Criterion included, the Flatline default of 5 seconds was determined to be sufficient for detecting flat channels and was not manipulated further. We evaluated the outputs from each criterion for bad channel detection relative to the manually selected channels by summing the number of false negatives and false positives for each file and calculating the overall accuracy rate across files for that set of automated parameters. False negatives refer to channels that were manually marked as bad but not flagged as bad by the pipeline. False positives refer to channels that were manually marked ‘good’ but were marked bad by the pipeline. An extra emphasis was placed on finding the settings with high accuracy that produced the lowest number of false positives in order to avoid getting rid of usable channels in the low-density dataset. HAPPILEE’s optimal settings produced 12 false negative and 5 false positive channels across all 19 files (228 total channels), with an overall accuracy rate of 95.5%.

**Table 1.** Performance of bad channel detection parameters tested on nineteen files from the example dataset. Pre-wav denotes that spectrum evaluation was run prior to wavelet thresholding and post-wav denotes that spectrum evaluation was run following wavelet thresholding.

| Bad Channel Parameters | Accuracy<br>(228 Total Channels) | False Positives<br>(193 Total Good Channels) | False Negatives<br>(35 Total Bad Channels) |
| --- | --- | --- | --- |
| Flatline 5; Channel Corr .7; Line Noise Ratio 2.5; Spectrum -2.75, 2.75 (pre-wav) | 95.5% | 5 | 12 |
| Flatline 5; Channel Corr .7; Line Noise Ratio 2; Spectrum -2.75, 2.75 (pre-wav) | 92.1% | 8 | 10 |
| Flatline 5; Channel Corr .7; Line Noise Ratio 2; Spectrum -2.75, 2.75 (pre-wav); Spectrum -2.5, 2.5 (post-wav) | 91.7% | 11 | 8 |
| Flatline 5; Channel Corr .7; Line Noise Ratio 2.5; Spectrum -2.75, 2.75 (pre-wav); Spectrum -2.5, 2.5 (post-wav) | 91.7% | 8 | 11 |
| Flatline 5; Channel Corr .7; Line Noise Ratio 2.5; Spectrum -2.75, 2.75 (pre-wav); Spectrum -2.4, 2.4 (post-wav) | 91.7% | 11 | 8 |
| Flatline 5; Channel Corr .7; Line Noise Ratio 2.85; Spectrum -2.75, 2.75 (pre-wav); Spectrum -2.5, 2.5 (post-wav) | 91.2% | 8 | 12 |
| Flatline 5; Channel Corr .7; Line Noise Ratio 2.5; Spectrum -2.5, 2.5 (pre-wav) | 90.8% | 10 | 11 |
| HAPPE 1.0 Legacy Detection | 85.1% | 0 | 34 |
| FASTER within MADE | 83.8% | 2 | 35 |

HAPPILEE combines EEGLab’s Clean Rawdata functions with power spectral evaluation steps as follows. HAPPILEE first runs the Clean Rawdata ‘Flatline Criterion,’ to detect bad channels with flat recording lengths longer than 5 seconds (indicating no data collected at that location). After flat channels have been removed, HAPPILEE uses Clean Rawdata’s ‘Line Noise Ratio Criterion’ with a threshold of 2.5 standard deviations (channels with line noise: neural data ratios greater than 2.5 standard deviations are marked as bad) and ‘Channel Correlation Criterion’ with a minimal acceptable correlation of 0.7 to detect additional bad channel cases. Finally, HAPPILEE includes a spectrum-based bad channel detection step following the Clean Rawdata functions. While the HAPPE 1.0 method of legacy detection proved to be insufficient for our low-density dataset (see Table 1), evaluating the joint probability of average log power from 1 to 100 Hz was useful for optimizing bad channel detection alongside Clean Rawdata. A spectrum evaluation step with thresholds of -2.75 and 2.75 was included to optimize bad channel detection accuracy. Thus, HAPPILEE achieves bad channel detection that is suitable for low density data and expands the classes of bad channels that can be detected relative to HAPPE 1.0 and MADE pipelines’ prior automated pipeline options.

### Artifact Rejection in Continuous Data

Raw EEG data may contain a number of artifacts (e.g. from participant motion, electromyogenic activity, eye movements/blinks) that must be removed from the signal during processing. Historically, artifact removal has been achieved via manual data inspection, where artifact-laden timepoints are deleted from the data. Automated artifact rejection approaches in high-density EEG pipelines to date have relied on independent component analysis (ICA) and wavelet thresholding methods instead as they can successfully reject artifact while retaining the entire length of the data file. ICA clusters data across electrodes into independent components that can segregate artifacts from neural data, while wavelet-thresholding parses data within frequency ranges using coefficients that can detect artifact fluctuations in either electrode data or independent components. ICA is included as an artifact rejection approach in many pipelines, including MADE and HAPPE 1.0 software. Wavelet thresholding was also implemented in HAPPE 1.0 as part of an initial wavelet-enhanced ICA (W-ICA) artifact rejection step. W-ICA entails first performing an ICA decomposition of the EEG signal into components to help cluster artifacts, after which all of the components’ timeseries are subjected to wavelet transform and thresholded to remove artifact before all of the components’ timeseries are translated back to EEG channel format (Castellanos and Makarov, 2006). Importantly, ICA is not an effective artifact rejection tool for low-density EEG configurations, as the number of channels determines the number of independent components to be generated through ICA, and many low-density configurations have too few channels to adequately segregate artifact from neural components. However, the wavelet thresholding approach (incorporated into the W-ICA step in HAPPE 1.0) provides an artifact rejection method that is suitable for low-density layouts, and preliminary work has found that wavelet thresholding outperforms ICA as an artifact removal approach on data with fourteen channels (Bajaj et al., 2020). Additionally, wavelet thresholding has been found to outperform other denoising methods that could apply to low-density data, including Empirical Mode Decomposition and Kalman filtering (Salis et al., 2013).

Wavelet thresholding also has several advantages relative to other artifact rejection approaches. To start, unlike ICA, the wavelet thresholding step of W-ICA does not have channel requirements (i.e. needing a specific number of channels) in order to run effectively. Additionally, wavelet thresholding is able to remove artifact without removing discrete timepoints, an issue that is inherent to manual artifact rejection where whole data segments are rejected. Furthermore, because wavelet decompositions are not dependent on channel number or large amounts of data (like in ICA), wavelet thresholding performance is consistent across datasets from different layout densities. Relatedly, wavelet thresholding can detect and remove isolated and non-stereotyped artifacts easily without removing neural data from other points in time or other channels. Wavelet thresholding also requires far less computation time than either ICA or manual inspection. All of these features make wavelet thresholding suitable for removing artifact from low-density EEG layouts.

Three approaches were taken to test and optimize automated wavelet thresholding-based artifact rejection in HAPPILEE using the BEIP dataset. Prior to artifact rejection testing, all files were initially filtered and subjected to line noise removal. The approaches are detailed below. The first approach (the clean vs. artifact approach) involved selecting two 30-second segments within each participant’s EEG file. The first 30-second segment was heavily artifact laden while the second 30-second segment was determined to be clean and without considerable artifact by experienced researchers. This approach facilitated testing whether artifact was effectively and accurately removed via wavelet thresholding. That is, optimal artifact rejection performance would be characterized by 1) substantial (artifact) signal removal for the artifact-laden 30 second file, indicating sensitivity to artifacts, and 2) minimal signal removal for the clean 30-second file from the same individual, indicating specificity in signal rejection constrained to artifact. High levels of signal rejection in the clean 30-second files would indicate unnecessary data loss, while low levels of signal rejection in the artifact-laden 30-second files could indicate insufficient performance. Additionally, to ensure that artifact removal was not biased to certain frequencies, data correlations pre- and post-processing were evaluated at key frequencies spanning all canonical frequency bands in the clean and artifact-laden segments. Examples of the clean and artifact signals within an individual (before and after wavelet thresholding) are provided in Figure 1.

**Figure 1.**
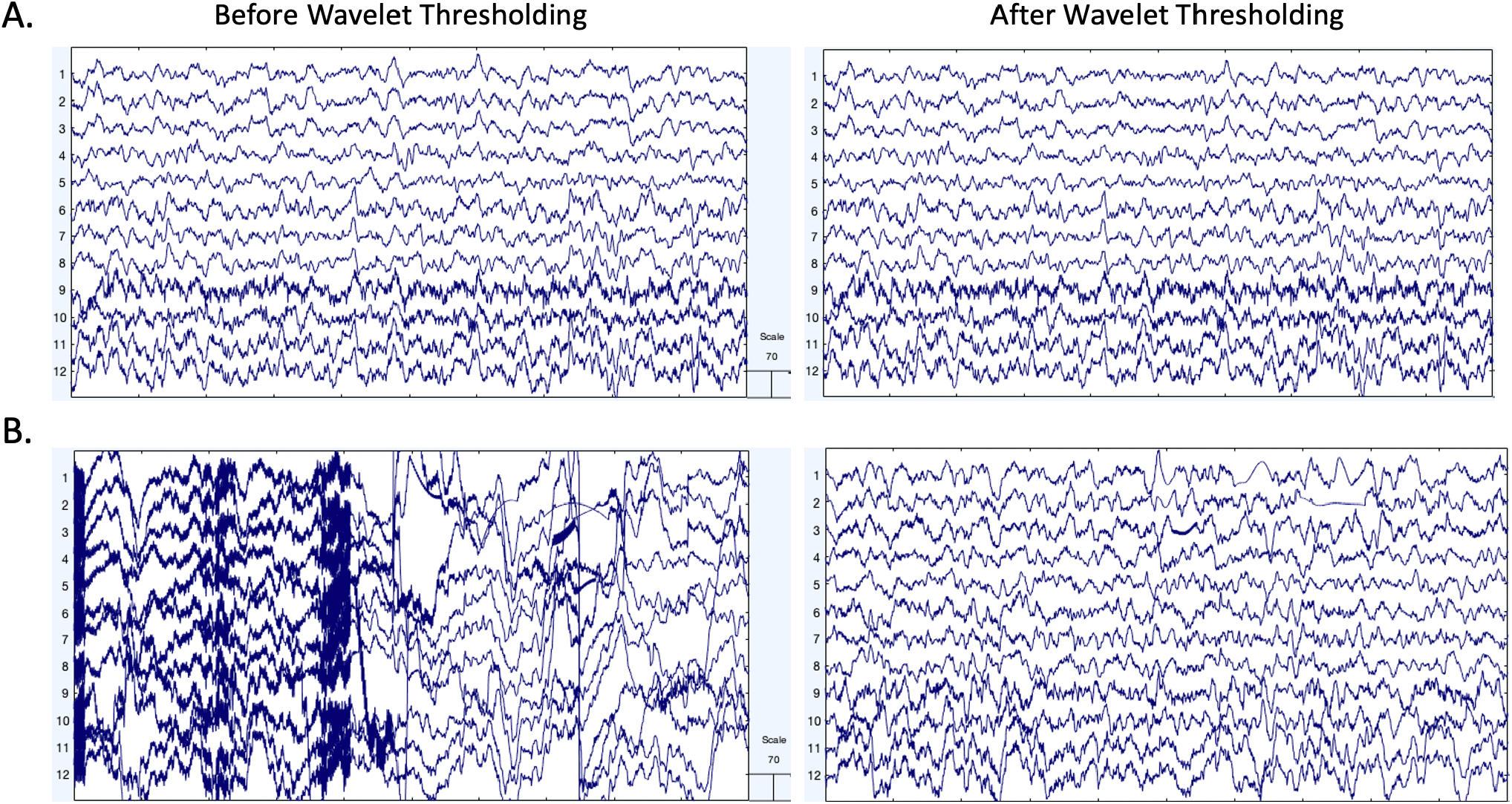
EEG signal before and after wavelet thresholding with the following parameters that were optimized for low density data: ‘Wavelet Family,’ ‘coif4;’ ‘Level of Decomposition,’ ‘10;’ ‘Noise Estimate,’ ‘Level Dependent;’ ‘Thresholding Method,’ ‘Bayes;’ ‘Threshold Rule,’ ‘Hard.’ Two files from the same participant in the example dataset are shown with 10 s of data extracted from the clean 30 second segment (A) and artifact-laden 30 second segment (B). The EEG signal before processing is shown in the left panel. The EEG signal after wavelet thresholding is shown in the right panel. All scales are in microvolts.

The second approach (artifact addition approach) used known artifact signals to establish how much artifact could be removed without distorting underlying neural signals during wavelet thresholding. In the absence of a ground-truth neural signal, we used the 30 second clean files from the first approach. We then isolated artifact timeseries by running ICA on the artifact laden 30-second files and selecting approximately 2 components per individual that were determined to be artifact with minimal neural data via visual inspection and automated classification through both the ICLabel and Multiple Artifact Rejection Algorithm options. We subsequently added those artifact timeseries on top of the clean 30-second data segment from the same individual. This artifact addition approach allowed us to evaluate how much of the added artifact was removed during wavelet thresholding by comparing the similarity of the artifact-added data post-wavelet thresholding to the post-wavelet thresholded clean 30-second segments (i.e. the original clean underlying signal) for the same participant (example from single participant in Figure 2).

**Figure 2.**
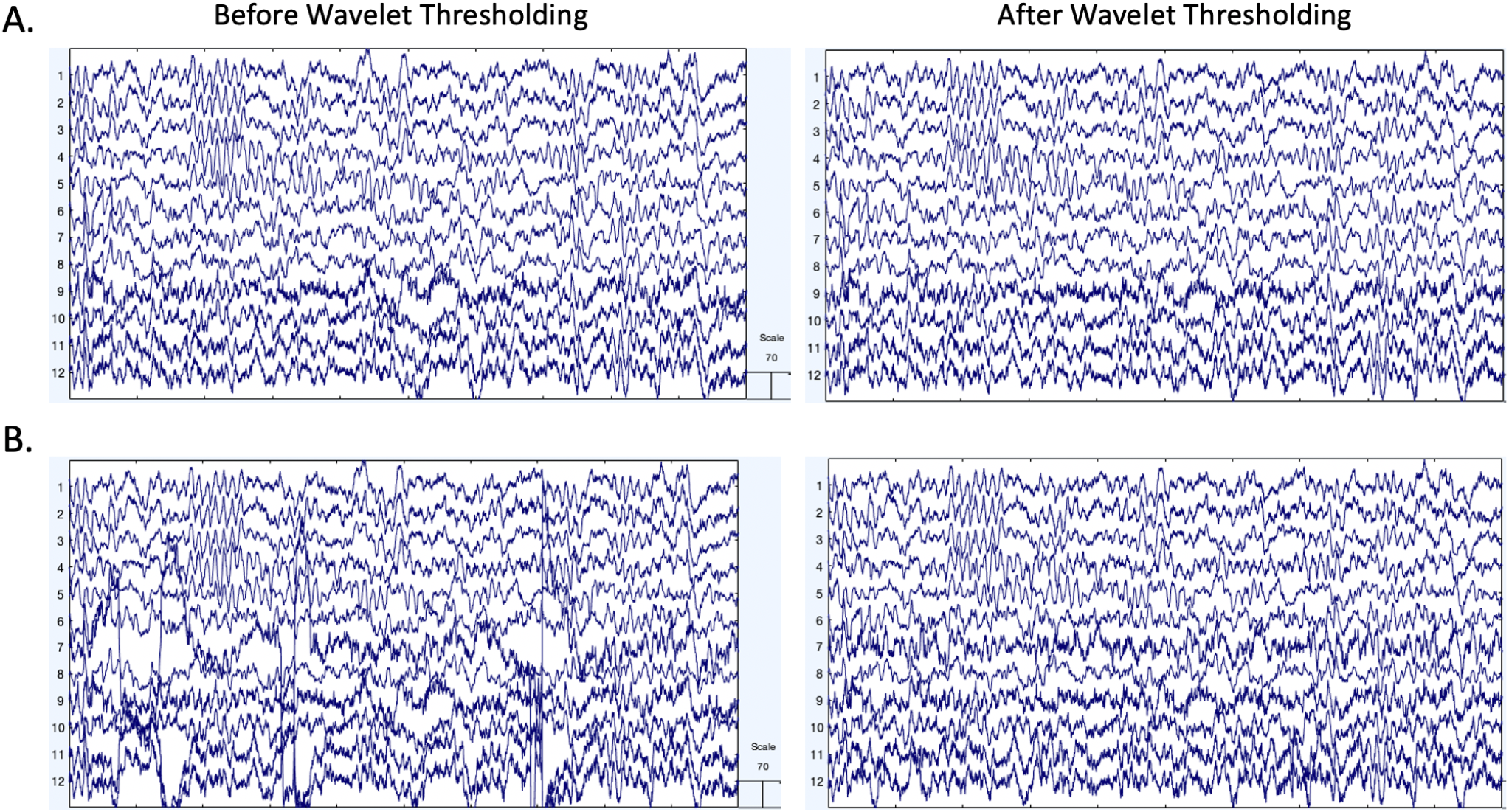
EEG signal before and after wavelet thresholding with the following parameters that were optimized for low density data: ‘Wavelet Family,’ ‘coif4;’ ‘Level of Decomposition,’ ‘10;’ ‘Noise Estimate,’ ‘Level Dependent;’ ‘Thresholding Method,’ ‘Bayes;’ ‘Threshold Rule,’ ‘Hard.’ Two files from the same participant in the example dataset are shown with 10 s of data extracted from the clean 30 second segment (A) and artifact-added 30 second segment (B). The EEG signal before processing is shown in the left panel. The EEG signal after wavelet thresholding is shown in the right panel. All scales are in microvolts.

For the third approach, we used simulated EEG data with artifact added to it in order to have a “ground truth” signal that we could attempt to recover with waveleting. To create the simulated EEG data, we used code from Bridwell et al. (2018). In short, the simulated EEG consisted of four signals. The four signals had distinct spatial patterns and frequency ranges (1.00-3.91Hz, 3.91-7.81Hz, 7.81-15.62Hz, and 15.62-31.25Hz). For a more thorough description on how the simulated signals were created, see Bridwell et al., 2018). After creating the simulated EEG data, we added blink and muscle artifact to the data to test various waveleting settings and wavelet thresholding in general. To get the blink artifact, we used a clear blink independent component (IC) from an adult participant (see Leach et al., 2020 for the specific study details). This IC was selected based on both an automated artifactual IC detection algorithm and visual inspection by two researchers with over five years of EEG and at least two years of ICA experience. For the muscle artifact, we pulled eight muscle ICs from the BEIP dataset used above in the artifact addition approach. After adding the artifact to the simulated EEG data, we did a 1Hz highpass and 35 Hz lowpass filter. Following this, we epoched the data into two-second epochs (50% overlap) to prepare the data for wavelet thresholding and/or artifact rejection. For artifact rejection, we used a -100 to 100 μV voltage threshold to identify bad epochs. We also required both frontal electrodes to exceed this threshold in order to classify an epoch as containing a blink. In order to test how well wavelet thresholding removes artifact without removing neural activity, we computed the power spectral density (PSD) on the original simulated signal (no artifact added) and compared that to the PSD of the signal after preprocessing with wavelet thresholding and without wavelet thresholding (i.e., the just artifact rejection condition). We further divided the wavelet thresholding condition into two more conditions that looked at how the preprocessed data might differ when waveleting is run before versus after artifact rejection.

This gave us three preprocessing conditions:

1. Artifact rejection only (no wavelet thresholding)
2. Wavelet thresholding followed by artifact rejection
3. Artifact rejection followed by wavelet thresholding

These three distinct approaches facilitated optimizing and evaluating the wavelet thresholding method for artifact removal in low-density data in multiple ways. Importantly, the wavelet thresholding approach broadly includes decomposing the EEG signal via a wavelet transform, determining a threshold value or values used to dissect data into the portion to be retained and the portion to be rejected, removal of the rejected data components, and reconstruction of the remaining signal. Each of these steps may be accomplished multiple ways across wavelet thresholding methods. Here, five key parameters in the wavelet thresholding process were manipulated and tested to optimize wavelet thresholding performance in this context, specifically: wavelet family, wavelet resolution (i.e. level of data decomposition), noise estimate method, thresholding level, and threshold rule. Each parameter was manipulated one at a time within a default set of wavelet thresholding parameters and tested using the clean vs. artifact approach and artifact injection approach in the BEIP dataset. The initial default set of wavelet thresholding parameters was chosen based on preliminary visual inspection of performance across a broader range of parameters prior to optimization and was as follows: ‘Wavelet Family,’ ‘coif5;’ ‘Level of Decomposition,’ ‘8;’ ‘Noise Estimate,’ ‘Level Dependent;’ ‘Thresholding Method,’ ‘Bayes;’ ‘Threshold Rule,’ ‘Soft.’ Subsequent optimization of each parameter is described in detail below.

#### Wavelet Family

The wavelet-thresholding method first subjects each electrode’s time series to wavelet transform by fitting a wavelet function to the data. The wavelet transform produces a series of coefficients to describe the EEG signal’s fluctuations across multiple frequency ranges. The wavelet function consists of both a wavelet family, dictated by the mother wavelet shape (e.g. the Coiflet mother wavelet is more symmetric than the Daubechies mother wavelet), and the wavelet order, which modifies the mother wavelet shape (see Figure 3; i.e. Coiflet order 4 wavelet has 8 vanishing moments in the function). To find the optimal wavelet function to carry out stationary wavelet transform, we tested the Coiflets family (orders 3, 4, and 5), the Daubechies family (orders 4 and 10), and the Symlets family (order 4). We selected these family/order combinations as they share shapes similar to those found in EEG signals and they are all orthogonal wavelet functions, which optimizes decomposition and reconstruction of the EEG signal from the wavelet transform (Strang and Nguyen, 1996). Moreover, prior literature indicates these wavelet families and specific orders have performed well on electrophysiological data (Al-Qazzaz et al., 2015; Alyasseri et al., 2017; Harender and Sharma, 2018; Lema-Condo et al., 2017; Nagabushanam et al., 2020). Using these wavelet families, we evaluated whether there was biased data rejection at any of the data frequencies in the clean 30 second segments by evaluating correlations between data pre-waveleting and post-waveleting at specific canonical frequencies. We also compared data rejection rates between the clean files and the 30 second artifact laden files. As a second analysis, we evaluated which wavelet family and order combination removed the most added artifact across frequencies (i.e. which wavelet facilitated the greatest correlation between the artifact added files and the clean files post-processing).

**Figure 3.**
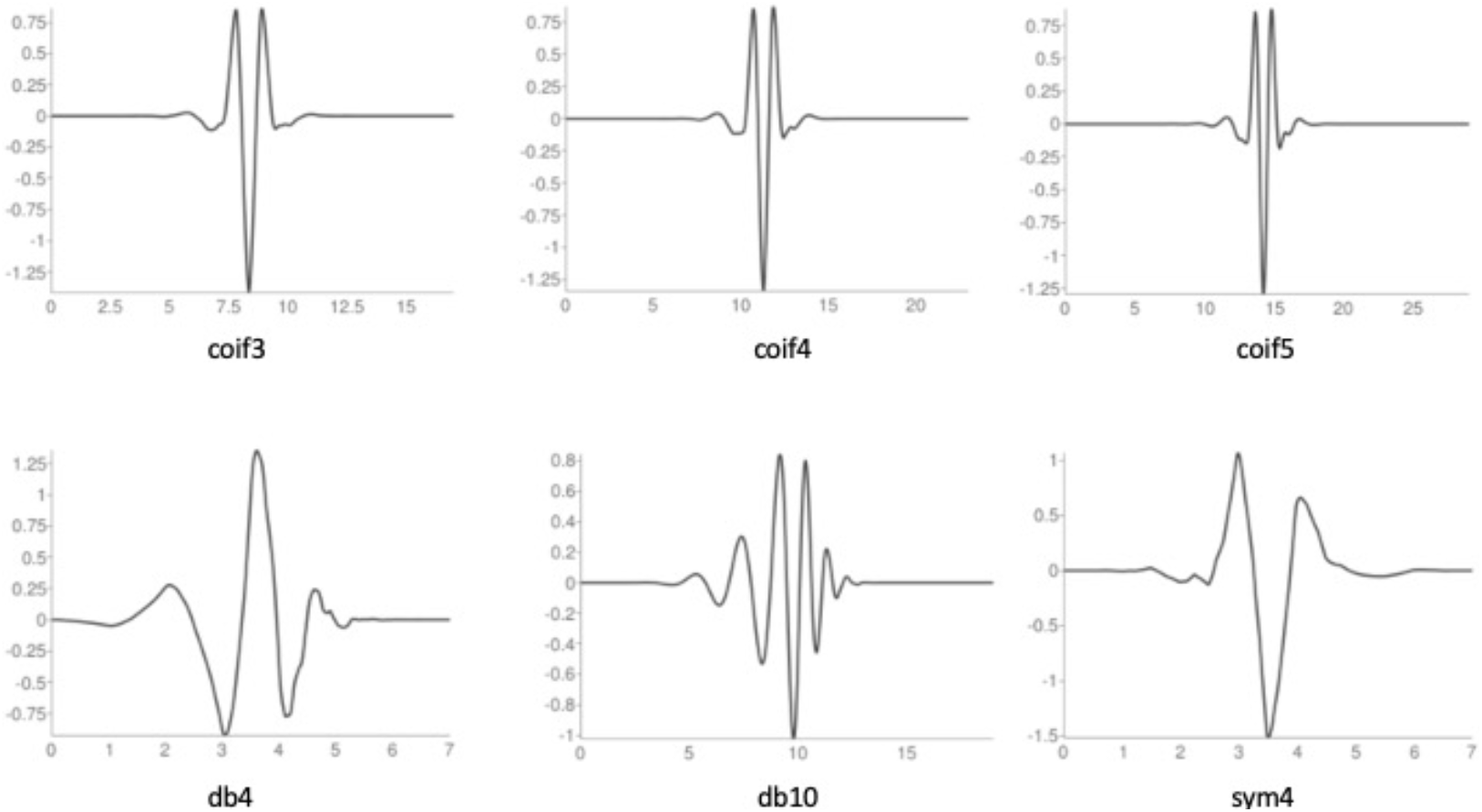
Visualization of the three wavelet families with selected orders tested to find the optimal wavelet function to carry out stationary wavelet transform on low density data.

After running the various wavelet family/order options, we found there were not meaningful differences between several wavelet family/order options (Tables 2 & 3). Specifically, performance did not differ between coif4, coif5, db4, and sym4 options across the clean vs. artifact-laden and artifact addition tests (e.g., correlation values in the artifact-addition tests were identical to the hundredths place). Coif4 was selected as the wavelet implemented in HAPPILEE as this wavelet/order also performed very well in data collected from a saline-based system (EGI), suggesting its performance may generalize more broadly than the other options tested here.

**Table 2:**
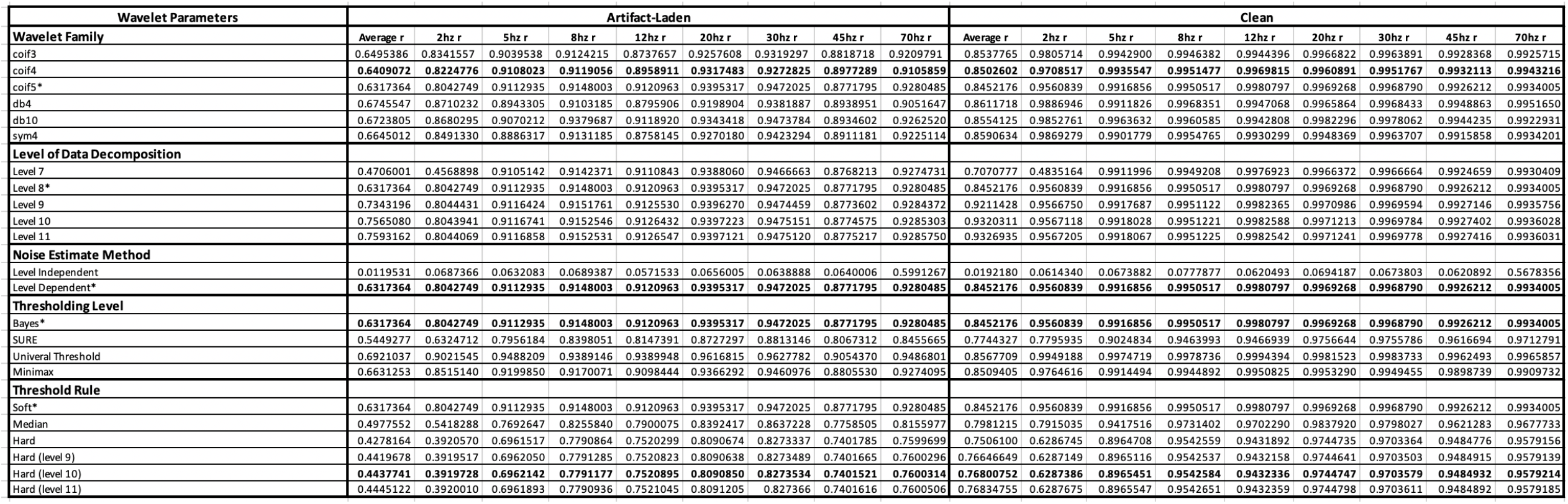
Artifact-added approach. Correlations between the EEG signal before wavelet thresholding and the EEG signal after wavelet thresholding for the wavelet parameters tested on the artifact-added 30 second segments. The r values of the wavelet parameters that are included in the final code are bolded. Asterisks denote default wavelet parameters.

#### Wavelet Resolution/Level of Data Decomposition

Following wavelet family/order selection, we manipulated the resolution of the wavelet that affects the level of data decomposition in wavelet thresholding. Specifically, this level of decomposition determines how fine-grained the frequency bands are in which data rejection occurs. Importantly, in the current code, the data sampling rate (not for example, the frequencies retained through initial filtering) determines which frequencies fall into different levels of decomposition. For example, the first level of decomposition for a file sampled at 1000 Hz (regardless of frequency filtering) would split data into two halves around 500 Hz. If that file had been resampled to 500 Hz prior to wavelet decomposition, the first level of decomposition would now split data into halves around 250 Hz. The default decomposition level splits data down to ≤1 Hz. We tested decomposition levels of ∼4 Hz, 2 Hz, and 1 Hz. The 4 Hz decomposition resulted in increased data removal in the lower frequencies relative to other frequencies of the clean data, resulting in data correlations pre-/post-thresholding of less than 0.5 (e.g., *r* = .48 at 2 Hz). This pattern indicated the need for further decomposition levels to avoid over-rejecting low frequency data that is not artifact-laden (low frequencies were rejected at similar rates in the artifact-laden data). In the clean vs. artifact files, we saw no difference in which data was rejected and retained when comparing levels 2 Hz and 1 Hz, though 1 Hz provides coverage down to the filtering cutoff. Therefore, HAPPILEE decomposes data into detail coefficients for frequencies above approximately 1 Hz to evaluate artifacts within each frequency range.

#### Noise (Artifact) Estimation Level

Once the level of decomposition is set, the noise estimate parameter is chosen to establish either a threshold for each level of decomposition (level dependent threshold) or establish a threshold that operates across all of the levels of decomposition (level independent threshold). The threshold(s) determine which wavelet coefficients describe data that is artifact-laden (i.e. coefficients describing larger amplitude changes, or noise) and will be removed from the EEG data during this artifact rejection step. Due to the unavailability of level dependent thresholding for the wavelet function used at the time of its conception, HAPPE 1.0 employed level independent thresholding (with a different threshold method and rule as well). We anticipated that level dependent thresholding would improve artifact detection specificity because of its ability to scale within each frequency range (e.g. artifacts in gamma frequencies have smaller amplitudes than artifacts in delta frequencies), rather than apply across all frequencies at once (which may over-penalize low-frequency clean EEG data that has higher amplitudes than higher-frequency clean EEG data). The default set of parameters was run with both level dependent and level independent thresholding and confirmed our prediction. With the improved thresholding method and rule included in the default parameters, the level independent threshold now heavily over-rejected the clean data (resulting in a correlation pre-/post-thresholding of *r* = .02), removing nearly all of the data (and a similar level of data rejection was observed in the artifact-laden data). (Note: this performance differs from HAPPE 1.0 due to the other wavelet thresholding parameter changes included in the new default settings of HAPPILEE, and does not reflect the functionality of wavelet-independent thresholding in HAPPE 1.0). There was no meaningful difference between level independent and level dependent thresholding in how much artifact was removed in the artifact-injection approach. This pattern of results suggested the level dependent threshold was just as effective at removing artifact as the level independent threshold without also removing underlying clean neural signal. As a result, a level dependent threshold was chosen in order to preserve data without compromising artifact rejection success.

#### Thresholding Method and Threshold Rule

Wavelet coefficients are then subjected to thresholding, (here, in a level-dependent way) such that coefficients with values smaller than a determined threshold for that level have their contribution to the data substantially suppressed (similar to Jansen, 2001; You and Chen, 2005). For EEG data, this effectively isolates the artifact signals within each frequency level (which are then subtracted out of the original EEG signal to clean it). A number of high performing options to determine the thresholds separating artifact from clean EEG have been established in the literature (Anumala and Kumar. Pullakura, 2018; Estrada et al., 2011; Geetha and Geethalakshmi, 2011; Geetha and Geethalakshmi, 2011; Guo et al., 2020; Jiang et al., 2007), specifically ‘Empirical Bayes’ (Johnstone and Silverman, 2004), ‘SURE’ (Donoho and Johnstone, 1995), ‘Universal’ (Donoho, 1995), and ‘Minimax’ (Donoho and Johnstone, 1998) thresholding approaches. HAPPE 1.0 originally included a Universal Threshold approach, but we aimed to examine these additional high-performing options as well to find the best fit for the current version of the pipeline. Relatedly, we evaluated available threshold rules for the various thresholding methods, specifically ‘Soft’ (Dai-Fei Guo et al., 2002; Donoho, 1995), ‘Median’ (Abramovich et al., 1998; Johnstone and Silverman, 2005), and ‘Hard’ (Dai-Fei Guo et al., 2002). Of note, not all thresholding methods are compatible with all thresholding rules. For example, the median rule is specific to the thresholding method ‘Bayes’ as it involves using the median posterior generated by the Bayesian algorithm. Soft and hard threshold rules are different in how they treat the coefficients near the threshold (in soft thresholding, these coefficients are shrunk while they are unaffected in hard thresholding). For thresholding options, we found minimal difference between options when tested on the artifact-added data, but we were able to eliminate ‘SURE’ due to over-rejecting data in the clean files (resulting in a correlation pre-/post-thresholding of *r* = .77). ‘Bayes’ narrowly outperformed ‘Minimax’ on the artifact-added data, but the reverse was true for our clean and artifact-laden data (Table 2). ‘Bayes’ was chosen as the thresholding method as it considers the uncertainty of the potential artifact and has been shown to result in more accurate denoising of signals generally. Moreover, the Bayes algorithm increases performance with increased data samples (as it can adjust its certainty estimates about artifact from more data), so performance on these 30-second files is a conservative reflection of this general performance. For threshold rule, on average, the median and hard thresholds removed more of the clean and artifact-laden file data, especially at the lower frequencies (e.g. at 2 Hz in clean data: soft threshold r = .84, median threshold r = 0.79, hard threshold r = 0.63, see Table 2). However, this effect was not present in the artifact-added data (Table 3), and visual inspection of data cleaned using soft vs. hard thresholds revealed that hard thresholds appeared to better remove artifact (see Figure 4). Therefore, this step was further explored using the simulated signals as described below and ultimately a Bayesian method with hard thresholding rule was implemented.

**Table 3:**
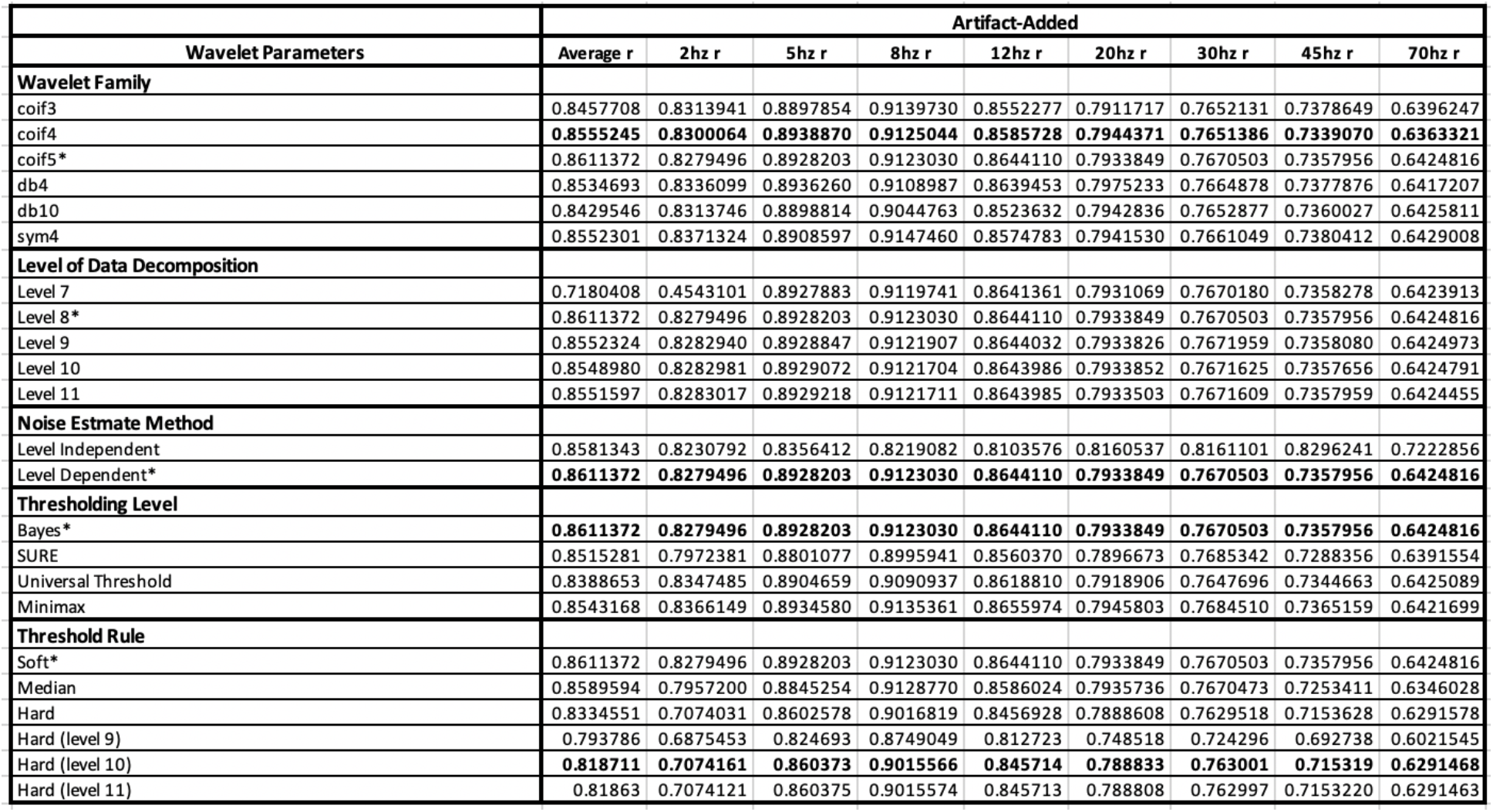
Artifact-added approach. Correlations between the EEG signal before wavelet thresholding and the EEG signal after wavelet thresholding for the wavelet parameters tested on the artifact-added 30 second segments. The *r* values of the wavelet parameters that are included in the final code are bolded. Asterisks denote default wavelet parameters.

|  | Artifact-Added |  |  |  |  |  |  |  |  |
| --- | --- | --- | --- | --- | --- | --- | --- | --- | --- |
| Wavelet Parameters | Average r | 2hz r | 5hz r | 8hz r | 12hz r | 20hz r | 30hz r | 45hz r | 70hz r |
| Wavelet Family |  |  |  |  |  |  |  |  |  |
| coif3 | 0.8457708 | 0.8313941 | 0.8897854 | 0.9139730 | 0.8552277 | 0.7911717 | 0.7652131 | 0.7378649 | 0.6396247 |
| coif4 | 0.8555245 | 0.8300064 | 0.8938870 | 0.9125044 | 0.8585728 | 0.7944371 | 0.7651386 | 0.7339070 | 0.6363321 |
| coif5* | 0.8611372 | 0.8279496 | 0.8928203 | 0.9123030 | 0.8644110 | 0.7933849 | 0.7670503 | 0.7357956 | 0.6424816 |
| db4 | 0.8534693 | 0.8336099 | 0.8936260 | 0.9108987 | 0.8639453 | 0.7975233 | 0.7664878 | 0.7377876 | 0.6417207 |
| db10 | 0.8429546 | 0.8313746 | 0.8898814 | 0.9044763 | 0.8523632 | 0.7942836 | 0.7652877 | 0.7360027 | 0.6425811 |
| sym4 | 0.8552301 | 0.8371324 | 0.8908597 | 0.9147460 | 0.8574783 | 0.7941530 | 0.7661049 | 0.7380412 | 0.6429008 |
| Level of Data Decomposition |  |  |  |  |  |  |  |  |  |
| Level 7 | 0.7180408 | 0.4543101 | 0.8927883 | 0.9119741 | 0.8641361 | 0.7931069 | 0.7670180 | 0.7358278 | 0.6423913 |
| Level 8* | 0.8611372 | 0.8279496 | 0.8928203 | 0.9123030 | 0.8644110 | 0.7933849 | 0.7670503 | 0.7357956 | 0.6424816 |
| Level 9 | 0.8552324 | 0.8282940 | 0.8928847 | 0.9121907 | 0.8644032 | 0.7933826 | 0.7671959 | 0.7358080 | 0.6424973 |
| Level 10 | 0.8548980 | 0.8282981 | 0.8929072 | 0.9121704 | 0.8643986 | 0.7933852 | 0.7671625 | 0.7357656 | 0.6424791 |
| Level 11 | 0.8551597 | 0.8283017 | 0.8929218 | 0.9121711 | 0.8643985 | 0.7933503 | 0.7671609 | 0.7357959 | 0.6424455 |
| Noise Estimate Method |  |  |  |  |  |  |  |  |  |
| Level Independent | 0.8581343 | 0.8230792 | 0.8356412 | 0.8219082 | 0.8103576 | 0.8160537 | 0.8161101 | 0.8296241 | 0.7222856 |
| Level Dependent* | 0.8611372 | 0.8279496 | 0.8928203 | 0.9123030 | 0.8644110 | 0.7933849 | 0.7670503 | 0.7357956 | 0.6424816 |
| Thresholding Level |  |  |  |  |  |  |  |  |  |
| Bayes* | 0.8611372 | 0.8279496 | 0.8928203 | 0.9123030 | 0.8644110 | 0.7933849 | 0.7670503 | 0.7357956 | 0.6424816 |
| SURE | 0.8515281 | 0.7972381 | 0.8801077 | 0.8995941 | 0.8560370 | 0.7896673 | 0.7685342 | 0.7288356 | 0.6391554 |
| Universal Threshold | 0.8388653 | 0.8347485 | 0.8904659 | 0.9090937 | 0.8618810 | 0.7918906 | 0.7647696 | 0.7344663 | 0.6425089 |
| Minimax | 0.8543168 | 0.8366149 | 0.8934580 | 0.9135361 | 0.8655974 | 0.7945803 | 0.7684510 | 0.7365159 | 0.6421699 |
| Threshold Rule |  |  |  |  |  |  |  |  |  |
| Soft* | 0.8611372 | 0.8279496 | 0.8928203 | 0.9123030 | 0.8644110 | 0.7933849 | 0.7670503 | 0.7357956 | 0.6424816 |
| Median | 0.8589594 | 0.7957200 | 0.8845254 | 0.9128770 | 0.8586024 | 0.7935736 | 0.7670473 | 0.7253411 | 0.6346028 |
| Hard | 0.8334551 | 0.7074031 | 0.8602578 | 0.9016819 | 0.8456928 | 0.7888608 | 0.7629518 | 0.7153628 | 0.6291578 |
| Hard (level 9) | 0.793786 | 0.6875453 | 0.824693 | 0.8749049 | 0.812723 | 0.748518 | 0.724296 | 0.692738 | 0.6021545 |
| Hard (level 10) | 0.818711 | 0.7074161 | 0.860373 | 0.9015566 | 0.845714 | 0.788833 | 0.763001 | 0.715319 | 0.6291468 |
| Hard (level 11) | 0.81863 | 0.7074121 | 0.860375 | 0.9015574 | 0.845713 | 0.788808 | 0.762997 | 0.7153220 | 0.6291463 |

**Figure 4.**
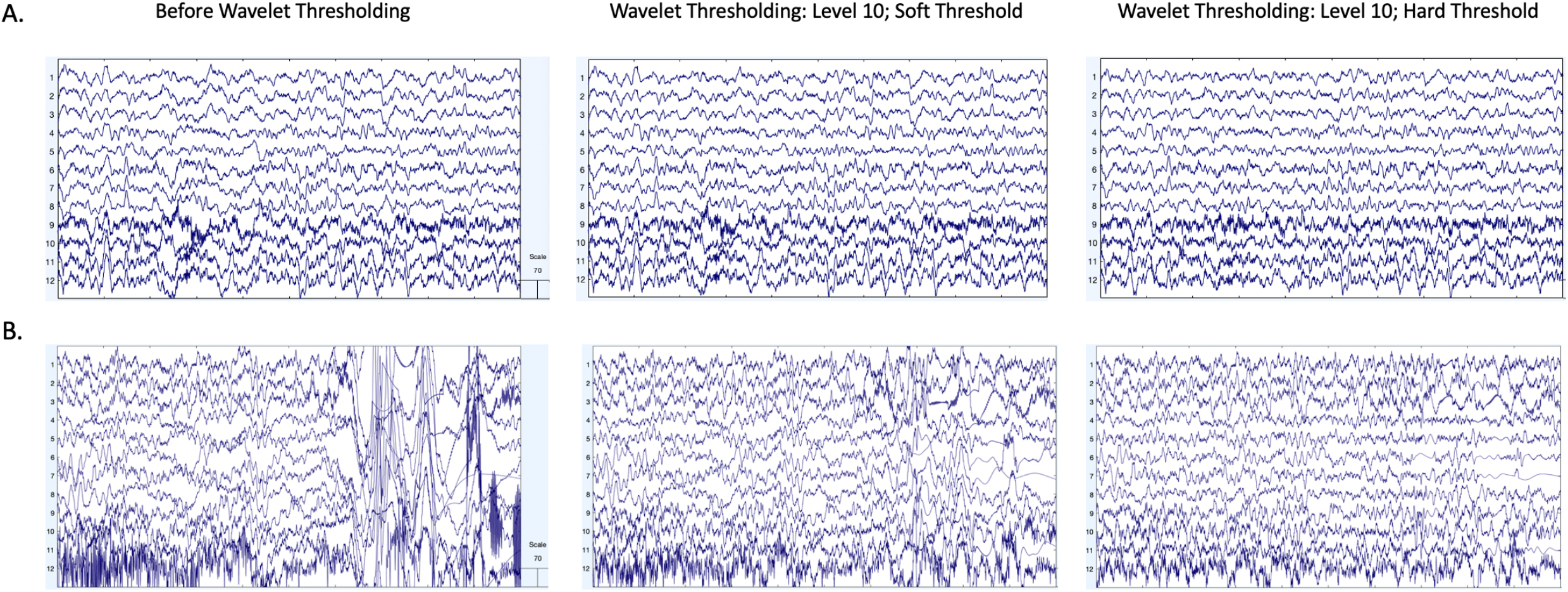
EEG signal before and after wavelet thresholding with the following parameters: ‘Wavelet Family,’ ‘coif5;’ ‘Level of Decomposition,’ ‘10;’ ‘Noise Estimate,’ ‘Level Dependent;’ ‘Thresholding Method,’ ‘Bayes;’ ‘Threshold Rule,’ ‘Soft’ (middle panel) and ‘Wavelet Family,’ ‘coif5;’ ‘Level of Decomposition,’ ‘10;’ ‘Noise Estimate,’ ‘Level Dependent;’ ‘Thresholding Method,’ ‘Bayes;’ ‘Threshold Rule,’ ‘Hard’ (right panel). Two files from the example dataset are shown with 10 s of data extracted from the clean 30 second segment (A) and artifact-laden 30 second segment (B). The EEG signal before processing is shown in the left panel. The EEG signal after wavelet thresholding with a soft threshold is shown in the middle panel. The EEG signal after wavelet thresholding with a hard threshold is shown in the right panel. All scales are in microvolts.

For the third approach with simulated data that had real artifact added to it, in terms of removing artifact while retaining the ground-truth underlying signal, wavelet thresholding performed better than just artifact rejection. Out of the nine wavelet thresholding function settings, we will highlight two settings that showed clear differences. First, the hard versus soft threshold setting better recovered the signal in lower frequency bands (Figure 5 top panel). Second, the level dependent versus level independent setting better recovered the signal in all frequency bands (Figure 5 bottom panel). Comparisons for the other seven settings showed only minor differences in their ability to remove artifact while retaining the original signal.

**Figure 5.**
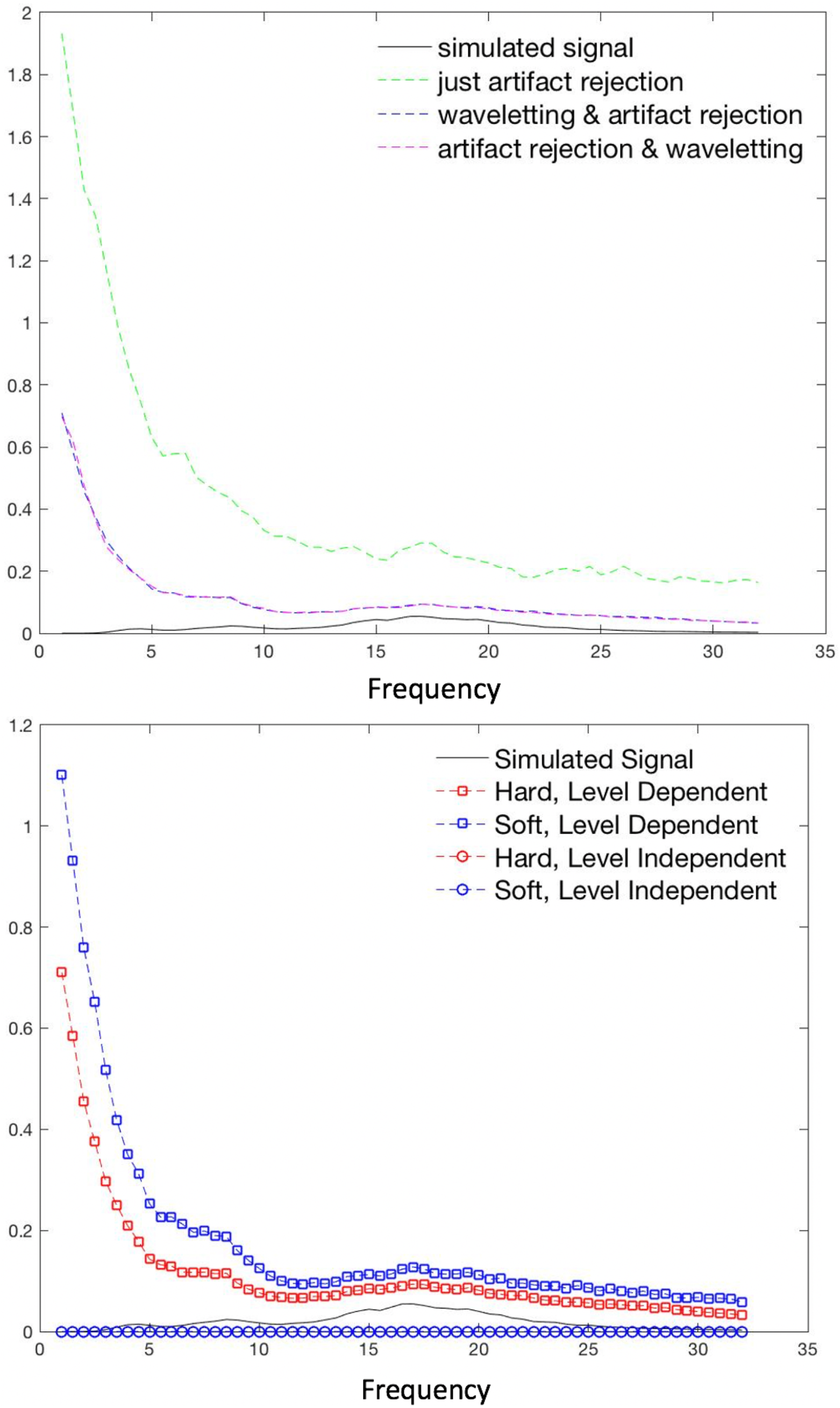
**(top panel)** EEG power spectral density after three conditions of pre-processing as compared to simulated signal. Y-axis is in microvolts. **(bottom panel)** EEG power spectral density after pre-processing with various wavelet thresholding parameters as compared to simulated signal. The two level-independent results are overlapping in this visual.

Finally, we examined whether the inclusion of wavelet-thresholding reduced the need to reject segments or trials of data downstream in the pipeline. To do so, we compared segment rejection rates in both the simulated and real EEG datasets. With respect to the simulated data, only wavelet thresholding approaches retained all trials (Table 4). As expected, when using artifact rejection, epochs exceeding the voltage threshold were removed, including some epochs with blinks, but not all. A total of 116 epochs were contaminated with blink artifact, but only 17 epochs were removed with voltage threshold artifact rejection when muscle artifact was included in the simulated data (in a separate test, not reported in detail here, we ran the data with just blink artifact included and 107/116 epochs with blinks were removed with voltage threshold artifact rejection). Visual inspection of the data suggested that some of the blinks in noisier epochs were retained because the noise stemming from muscle activity included negative deflections that lowered the amplitude of the blinks, which caused the blink artifacts to fall within our acceptable voltage threshold range. If, however, only one frontal electrode needed to exceed our voltage threshold, 90/116 trials were correctly marked for rejection because of blink deflections. Wavelet thresholding appears (based on visual inspection) to have removed these blinks from the signal. However, some muscle activity still remained in the simulated data in all preprocessing conditions.

**Table 4.** Performance of three preprocessing sequences on simulated data.

|  | Artifact Rejection | Waveleting followed by artifact rejection | Artifact rejection followed by waveleting |
| --- | --- | --- | --- |
| Trials retained | 269 | 286 | 269 |
| Proportion of trials retained | 94% | 100% | 94% |

To evaluate segment retention in the real EEG data, manual segment rejection rates in 14 files of the BEIP dataset that were processed with and without waveleting on the data (mean number of segments before rejection = 105.6). The mean number of segments retained after manual rejection on post-waveleted data (85.7 segments) was significantly higher than the mean number of segments retained without wavelet thresholding (62.6 segments; t(18) = -9.07, p = 0.00000004). That is, wavelet-thresholding improved segment retention by 37% relative to traditional manual editing approaches, further evidence in support of its use in preprocessing EEG data.

The final wavelet-thresholding parameters implemented in HAPPILEE are as follows: ‘Wavelet Family,’ ‘coif4;’ ‘Level of Decomposition,’ ‘10;’ ‘Noise Estimate,’ ‘Level Dependent;’ ‘Thresholding Method,’ ‘Bayes;’ ‘Threshold Rule,’ ‘Hard’.

### Segmentation (Optional)

HAPPILEE includes an optional data segmentation step along with several additional artifact rejection steps to further optimize processing. For low density data with event markers (e.g., event-related EEG data or ERP designs), data can be segmented around events as specified by user inputs (ERP-processing is supported, including baseline and timing offset correction). For data without event markers (e.g., resting-state EEG), regularly marked segments of any duration specified by the user are generated for the duration of the recording (e.g. 1-second segments).

Users with data files where segment rejection would lead to an unacceptably low remaining number of segments for analysis may choose an optional post-segmentation step involving the interpolation of data within individual segments for channels determined to be artifact-contaminated during that segment, as implemented by FASTER software (Nolan et al., 2010). Each channel in each segment is evaluated on the four FASTER criteria (variance, median gradient, amplitude range, and deviation from mean amplitude), and the Z score (a measure of standard deviation from the mean) for each channel in that segment is generated for each of the four metrics. Any channels with one or more Z scores that are greater than 3 standard deviations from the mean for an individual segment are marked bad for that segment. These criteria may identify segments with residual artifacts. Subsequently, for each segment, the channels flagged as bad in that segment have their data interpolated with spherical splines, as in FASTER. This allows users to maintain the maximum number of available segments, while still maximizing artifact rejection within individual segments. However, we caution users from implementing this option in cases where channels are distributed with significant distance between them as the interpolation process would pull data from distal channels that does not reflect the appropriate activity profile for that scalp space.

The majority of users, including those who wish to avoid interpolating data within individual segments, may instead choose to reject segments that are determined to still be artifact-contaminated. HAPPILEE includes three segment rejection options. Criteria for rejection include a choice of joint-probability criteria, amplitude-based criteria, or a combination of joint-probability criteria with amplitude-based criteria. Joint-probability criteria considers how likely a segment’s activity is given the activity of other segments for that same channel, as well as other channels’ activity for the same segment. The assumption is that artifact segments should be the rare segments with activity several standard deviations apart relative to the rest of the data.

Amplitude-based criteria sets a minimum and maximum signal amplitude as the artifact threshold, with segments being removed when their amplitude falls on either side of this threshold. HAPPILEE allows the user to specify their minimum and maximum allowable amplitudes. Users may also specify a combined approach with joint-probably criteria and amplitude-based criteria that removes outlier segments that fail either standard deviations or the signal amplitude criteria.

To test the efficacy of the three segment rejection options and determine the optimal criterion values for the rejection of segments in low density EEG data, we compared a series of ten automated options to a set of manually rejected segments for fourteen files in the BEIP dataset. We manipulated the standard deviation values for joint-probability rejection, the amplitude values for amplitude-based rejection, and tested combinations of different joint-probability standard deviations with amplitude criteria. The number of segments rejected for each of these automated rejection approaches was compared to the segments rejected via manual inspection as the gold standard approach using paired t-tests (Table 5). Amplitude-based rejection alone did not sufficiently match the manual rejection rates. However, the number of segments rejected using joint-probability criteria of 2 standard deviations alone or in combination with amplitude criteria (here, -150 and 150 microvolts) were not significantly different from the number of segments rejected manually (both p > 0.1).

**Table 5.**
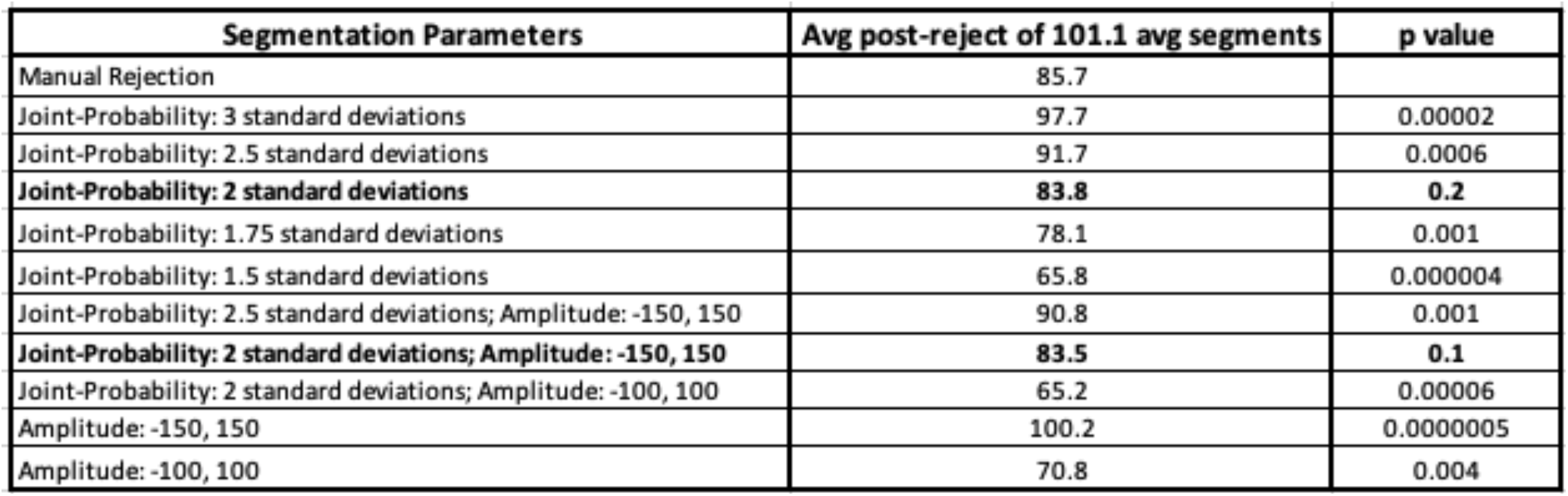
Comparison of various segment rejection options tested on nineteen files from the example dataset. P values are calculated from t-tests comparing the number of remaining segments for each parameter following automated rejection to the number of segments remaining following manual rejection.

Segment rejection performance was further evaluated for these two approaches by comparing the identity of segments rejected via the automated approach to the manually rejected segments by summing the number of false negatives and false positives for each file and calculating the overall accuracy rate across files compared to the manual rejection classification (i.e. did HAPPILEE reject the same segments that were rejected manually). The joint probability criterion alone (using 2 standard deviations) achieved the higher accuracy rate of 91.2% across all files but joint-probability with amplitude also did well (91.0% accuracy) (see Table 6). HAPILEE therefore includes three segment rejection options with the following recommendations. For data with sufficient channels (e.g. here 12 was sufficient but we did not test performance on sparser configurations), segment rejection via joint-probability criteria is recommended. This setting is also recommended for users combining data collected across different systems or ages where the overall signal amplitude may differ across files. Users performing analyses in the time domain as for ERP paradigms may opt to include amplitude-based criteria. For users with very low-density configurations where the joint-probability criteria relying on standard deviations may not perform as well as it did here, amplitude-only criteria may be used for segment rejection.

**Table 6.** Performance of segment rejection parameters tested on nineteen files from the example dataset.

| Segmentation Parameters | Accuracy | False Positive Rate | False Negative Rate |
| --- | --- | --- | --- |
| Joint-Probability: 2 standard deviations | 91.2% | 6.5% | 15.9% |
| Joint-Probability: 2 standard deviations; Amplitude: -150, 150 | 91.0% | 6.9% | 15.5% |

For all HAPPILEE runs that included bad channel detection, channels marked as bad are now subject to spherical interpolation (with Legendre polynomials up to the 7th order) to repopulate their signal. The identity of all interpolated channels, if any, for a file are recorded in the HAPPILEE processing report for users who wish to monitor the percentage or identity of interpolated channels in their datasets before further analysis. This interpolation step is available for all files formats except for .mat formats without channel locations as the interpolation step requires electrode location information to interpolate appropriately spatially.

### Re-referencing (Optional)

HAPPILEE offers users the choice to re-reference the EEG data. If re-referencing, the user may specify either re-referencing using an average across all channels (i.e., average re-reference) or using a channel subset of one or multiple channels. For both re-referencing options, only channels within the user-specified channel subset selected for HAPPILEE processing can be used for re-referencing. Re-referencing also reduces artifact signals that exist consistently across electrodes, including residual line-noise. During re-referencing, if there is a prior reference channel (e.g. an online reference channel), that channel’s data is recovered and included in the re-referenced dataset. For example, EGI data is typically online-referenced to channel CZ. In this example, users could now recover data at channel CZ by re-referencing to any other channel or channels in this step.

An example file pre- and post-processing with HAPPILEE is shown in Figure 6 to demonstrate the effectiveness of the pipeline. Clean 30-second segment of data pre- and post-processing is shown as well as an artifact-laden 30-second segment of data pre- and post-processing. The clean and artifact-laden power spectra are much more similar post-processing compared to pre-processing.

**Figure 6.**
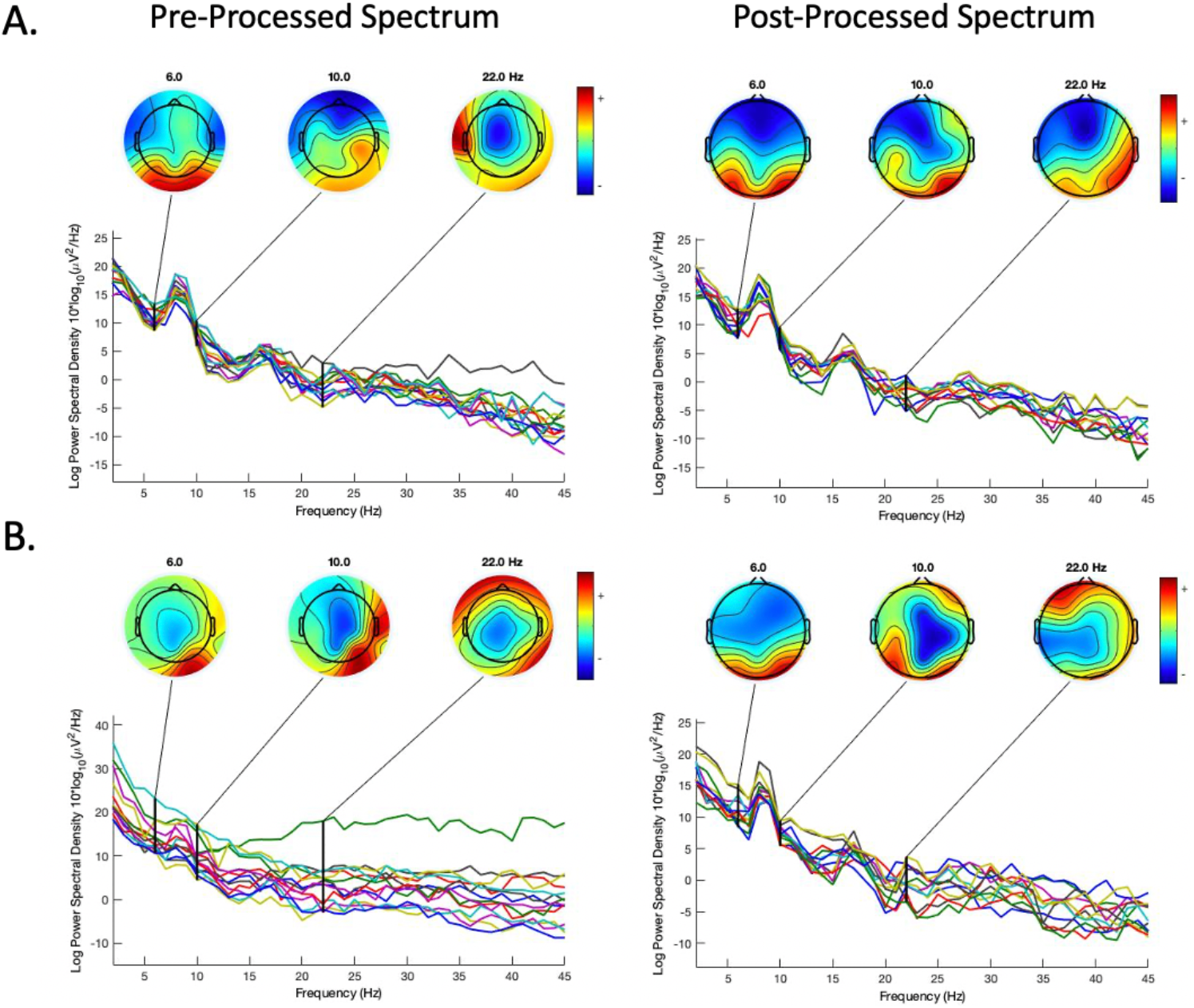
EEG spectrum before and after complete processing through HAPPILEE with the final optimizations for each step. Two files from the same participant in the example dataset are shown from the clean 30 second segment (A) and artifact-laden 30 second segment (B). The EEG spectrum before processing is shown in the left panel. The EEG spectrum after processing is shown in the right panel.

### HAPPILEE Outputs

HAPPILEE outputs include the processed EEG or ERP data and the HAPPILEE processing reports. These outputs are generated in several folders that are located within the user-specified folder of files for processing. EEG files are saved out after several intermediate processing steps so that users can explore in-depth and visualize how those steps affected the EEG signal in their own datasets. The intermediate files are separated into folders based on the level of processing performed on the data and include: 1) data after filtering to 100 Hz and line-noise reduction, 2) data post-bad channel rejection (if selected), and 3) post-wavelet-thresholded data. If segmenting is enabled, HAPPILEE outputs one to two additional intermediate files: 5) post-segmented EEG data (always) and 6) interpolated data (if bad data interpolation is enabled). If segment rejection is selected, HAPPILEE saves the data post-segment rejection as well.

HAPPILEE outputs fully processed files that are suitable inputs for further analyses in one of several formats, selected by the user at the start of the HAPPILEE run, to increase compatibility with other software for data visualizations or statistical analyses. Options include mat, .set, and .txt formats. Alongside the fully processed data, HAPPILEE also outputs the HAPPE Data Quality Assessment Report and the HAPPE Pipeline Quality Assessment Report, each described in detail below. Finally, if HAPPILEE is run in the semi-automated setting, the software generates an image for each file containing the fully processed data’s power spectrum.

### HAPPILEE Data Quality Assessment Report

HAPPILEE generates a report table of descriptive statistics and data metrics for each EEG file in the batch in a single spreadsheet to aid in quickly and effectively evaluating data quality across participants within or across studies. The report table with all these metrics is provided as a .csv file in the “quality_assessment_outputs” folder generated during HAPPILEE. We describe each of these metrics below to facilitate their use to determine and report data quality.

#### File Length in Seconds

HAPPILEE outputs the length, in seconds, of each file prior to processing.

#### Number of Segments Before Segment Rejection and Number of Segments Post Segment Rejection

HAPPILEE reports the number of segments before segment rejection and post segment rejection. If segment rejection is not enabled, these numbers are identical. If theuser enabled segment rejection in HAPPILEE, they may evaluate the number of data segments remaining post-rejection for each file to identify any files that cannot contribute enough clean data to be included in further analyses (user discretion). The user may also easily tabulate the descriptive statistics for remaining segments to report in their manuscript’s Methods section (e.g., the mean and standard deviation of the number of usable data segments per file in their study).

#### Percent Good Channels Selected and Interpolated Channel IDs

The percentage of channels contributing un-interpolated data (“good channels”) and the identity of interpolated channels are provided. Users wishing to limit the amount of interpolated data in further analyses can easily identify files for removal using these two metrics.

#### ICA-Related Metrics

As HAPPILEE does not perform ICA on low-density EEG data, the metrics measuring ICA performance in HAPPE 2.0 are assigned “NA.”

#### Channels Interpolated for Each Segment

If the user selected the Data Interpolation within Segments option of the additional segmenting options, HAPPILEE will output a list of segments and the channels interpolated within each segment for each file. Otherwise, it will output “N/A.” Users wishing to limit the amount of interpolated data in further analyses can easily identify files for removal using this metric.

### HAPPE Pipeline Quality Assessment Report

For each run, HAPPILEE additionally generates a report table of descriptive statistics and data metrics for each EEG file in the batch in a single spreadsheet to aid in quickly and effectively evaluating how well the *pipeline* performed across participants within or across studies. Note that these metrics may also be reported in manuscript methods sections as indicators of how data manipulations changed the signal during preprocessing. The report table with all these metrics is provided as a .csv file in the “quality_assessment_outputs” folder generated during HAPPILEE processing.

#### r pre/post linenoise removal

HAPPILEE automatically outputs cross-correlation values at and near the specified line noise frequency (correlation between data at each frequency before and after line noise removal). These cross-correlation values can be used to evaluate the performance of line noise removal, as the correlation pre- and post-line noise removal should be lower at the specified frequency, but not at the surrounding frequencies beyond 1 to 2 Hz. HAPPILEE will automatically adjust which frequencies are reported depending on the user-identified line noise frequency. This metric can also be used to detect changes in how much line noise is present during the recordings (e.g. if generally cross-correlation values are high when study protocol is followed, indicating low line-noise removal from the data, but a staff member forgets to remove their cell phone from the recording booth for several testing sessions, the degree of line noise removal for those files summarized by this metric could be used as a flag to check in on site compliance with acquisition protocols).

#### r pre/post wav-threshold

HAPPILEE automatically outputs the cross-correlation values before and after wavelet thresholding across all frequencies and specifically at 0.5 Hz, 1 Hz, 2 Hz, 5 Hz, 8 Hz, 12 Hz, 20 Hz, 30 Hz, 45 Hz, and 70 Hz. These specific frequencies were selected to cover all canonical frequency bands across the lifespan from delta through high-gamma as well as the low-frequencies retained in ERP analyses. These cross-correlation values can be used to evaluate the performance of waveleting on the data for each file. For example, if cross-correlation values are below 0.2 for all participants in the sample, the wavelet thresholding step has not progressed as intended (users are advised to first check their sampling rate in this case and visualize several raw data files). Note that this measure may also be used to exclude individual files from further analysis based on dramatic signal change during waveleting (indicating high degree of artifact), for example if the 8 Hz or all-data cross-correlations are below some threshold set by the user (e.g., 3 standard deviations from the median or mean, r values below 0.2).

Through these quality assessment reports, HAPPILEE aims to provide a rich, quantifiable, yet easily accessible way to effectively evaluate data quality for even very large datasets in the context of automated processing. Visual examination of each file is not required, although it is available. We also hope to encourage more rigorous reporting of data quality metrics in manuscripts by providing these outputs already tabulated and easily transformed into descriptive statistics for inclusion in reports. Users may also wish to include one or several of these metrics as continuous nuisance covariates in statistical analyses to better account for differences in data quality between files or verify whether there are statistically significant differences in data quality post-processing between study groups of interest.

Several metrics may also be useful in evaluating study progress remotely to efficiently track the integrity of the system and data collection protocols. For example, the r pre/post linenoise removal metric may indicate environmental or protocol deviations that cause significant increases in line noise in the data, and the Percent Good Channels Selected and Interpolated Channel ID metrics can be used to track whether the net/cap is being applied and checked for signal quality prior to paradigms or whether a channel (or channels) is in need of repair. For example, if the T6 electrode starts consistently returning bad data for a specific net/cap, it may need to be examined for repair. For further guidance about using the processing report metrics to evaluate data, users may consult the User Guide distributed with HAPPILEE software.

### Conclusion

The field has been rapidly moving toward the use of automated EEG preprocessing pipelines that make use of contemporary artifact-rejection approaches like wavelet-thresholding and ICA as effective, efficient, standardized alternatives to subjective and labor-intensive manual preprocessing. Here we provide a solution suitable for low-density layouts down to single channel EEG with the current automated pipeline, HAPILEE (see Figure 7 for final pipeline schematic). HAPILEE supports processing resting-state and task-related EEG, as well as ERP data. HAPILEE is suitable for configurations with any number of channels, though it may perform best on data with 1 to 32 channels (those with higher-density configurations may consider pipelines optimized for high-density data, including the companion HAPPE pipeline or HAPPE+ER pipeline within HAPPE 2.0 software).

**Figure 7.**
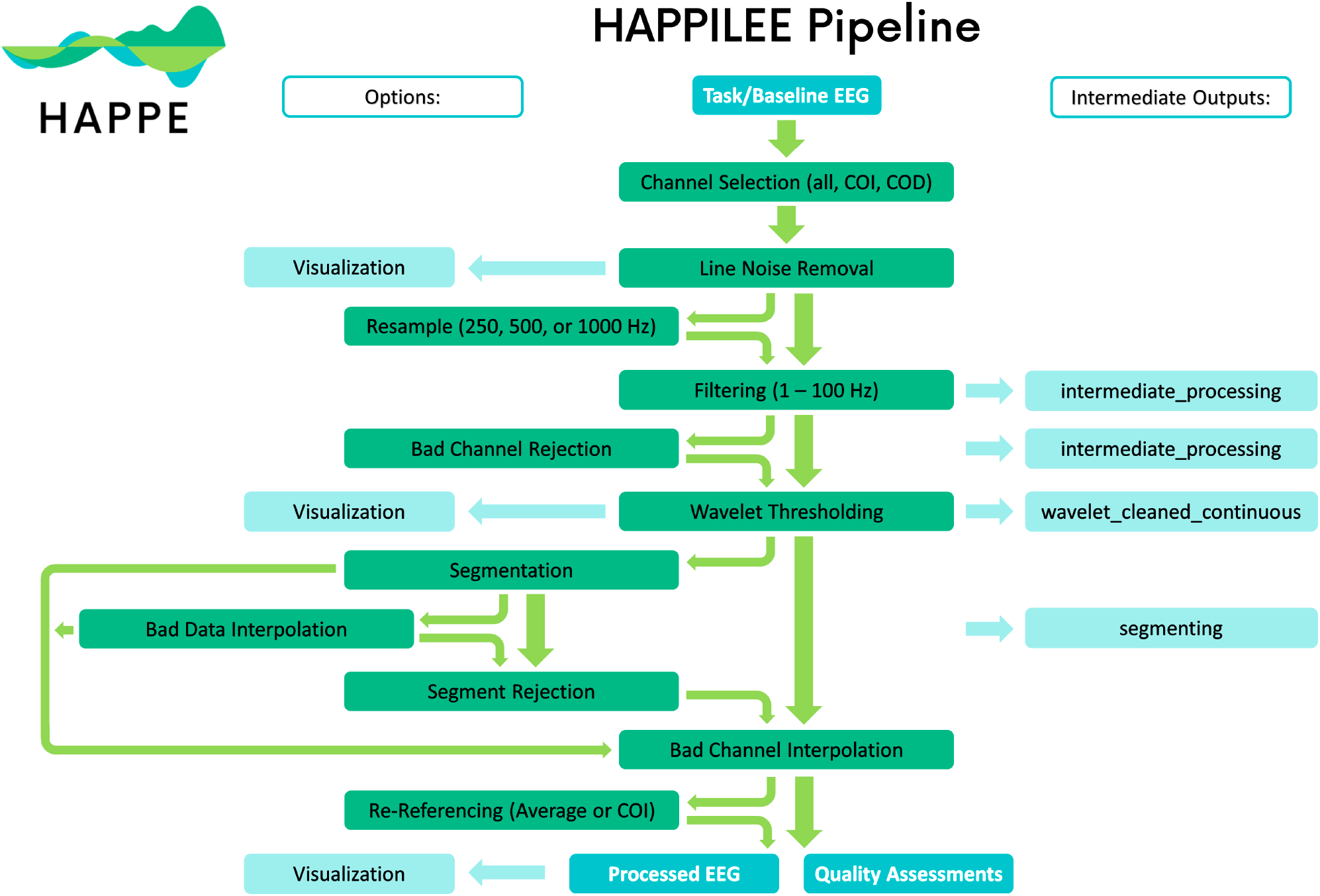
Schematic illustrating the HAPPILEE pipeline’s processing steps. The intermediate output EEG files are indicated by the suffix added after that specific processing step in the light blue boxes. The user options for resampling, segmentation, bad data interpolation, segment rejection, and re-referencing steps and visualizing several steps in HAPPILEE with the semi-automated setting are also indicated.

There are several limitations to HAPPLIEE that should also be considered. First, bad channel detection was tested on a dataset with twelve channels. As a result, those working with layouts with substantially fewer electrodes may consider verifying for themselves that the default settings work sufficiently well for their datasets. Alternatively, the bad channel detection step is optional, so if it is unsuitable or is not desired for a dataset, the user may opt-out of this step of the pipeline. Furthermore, the appropriate amplitude threshold for performing segment rejection by amplitude will vary across datasets collected on different ages or systems and should be verified through visual inspection of several files (via running HAPPILEE in the semi-automated setting with visualizations). Lastly, HAPPILEE was optimized using a single system (James Long) given the data available to the researchers, but others should independently verify performance on the other systems compatible with HAPPILEE.

The HAPPILEE pipeline is freely available as part of the HAPPE software (first released with HAPPE version 2.0), covered under the terms of the GNU General Public License (version 3) (Free Software Foundation, 2007). HAPILEE’s sequence of processing steps are automatically triggered within HAPPE 2.0+ software when the user indicates they have data with fewer than 32 channels. HAPPILEE may be accessed at: https://github.com/PINE-Lab/HAPPE. The subset of BEIP EEG data used to optimize the HAPILEE pipeline, including the files used in the clean vs. artifact and artifact addition approaches are publicly available at Lopez et al. on Zenodo. To facilitate use of HAPPILEE, a user guide is also included in the software download.

